# Nanoscale details of mitochondrial fission revealed by cryo-electron tomography

**DOI:** 10.1101/2021.12.13.472487

**Authors:** Shrawan Kumar Mageswaran, Danielle Ann Grotjahn, Xiangrui Zeng, Benjamin Asher Barad, Michaela Medina, My Hanh Hoang, Megan J Dobro, Yi-Wei Chang, Min Xu, Wei Yuan Yang, Grant J. Jensen

## Abstract

Mitochondrial fission is required for proper segregation during cell division, quality control, and cellular homeostasis (metabolism and energy production). Despite its importance, models of the process remain speculative. Here we apply cryogenic electron tomography to image the nanoscale architecture of mitochondrial fission in mammalian cells. We find that constriction of the inner and outer membranes is coordinated, suggesting that force on both membranes is applied externally. While we observe ER at constriction sites, it did not encircle constrictions. Instead, we find long bundles of both unbranched actin and septin filaments enriched at constrictions. Actin bundles align with the central region of division bridges and septin bundles with the necks on either side. Septin bundles appear to guide microtubules to constriction sites, suggesting, along with autolysosomes observed in the vicinity, a pathway for mitophagy. Together, our results rule out several existing models for mitochondrial fission and provide empirical parameters to inform the development of realistic coarse-grained models in the future.

## Introduction

Mitochondria are double-membraned, semi-autonomous organelles that perform critical functions in eukaryotic cells including ATP generation via the citric acid (Krebs) cycle and oxidative phosphorylation, biogenesis of amino acids, regulation of ion homeostasis, and initiation of apoptosis (Osellame et al., 2012). The dynamic balance between fission (one mitochondrion splitting into two) and fusion (two mitochondria joining into one) controls their shape, size, number and cellular distribution, which in turn affect their ability to perform their essential functions (Chan, 2006; Detmer and Chan, 2007; Friedman and Nunnari, 2014). Mitochondrial fragmentation via fission prior to cell division ensures the proper inheritance of the organelle by daughter cells (Mishra and Chan, 2014), and damaged regions of mitochondria are excised by fission to be degraded via mitophagy – targeted degradation of mitochondria by autophagosomes (Palikaras et al., 2018). Disruptions to fission are associated with etiologically diverse diseases including neurodegenerative disorders (Nunnari and Suomalainen, 2012; Serasinghe and Chipuk, 2017).

Much of the molecular machinery responsible for mitochondrial fission has been identified, but the structural mechanisms underlying the process are not well understood (Pagliuso et al., 2018; Tilokani et al., 2018). For instance, fission requires the constriction and scission of two separate membranes, the inner mitochondrial membrane (IMM) and the outer mitochondrial membrane (OMM), but the extent to which constriction of these two membranes is coordinated remains unclear. In mammalian cells, several cytosolic factors participate in fission, including the endoplasmic reticulum (ER) (Friedman et al., 2011), dynamin-related GTPases like Drp1 (Smirnova et al., 2001) and Dyn2 (Lee et al., 2016), cytoskeletal filaments such as actin (Korobova et al., 2013) and septin (Pagliuso et al., 2016), and the motor protein myosin II (Korobova et al., 2014; Yang and Svitkina, 2019). ER and actin filaments are thought to initiate mitochondrial constrictions and several models for their force generation have been proposed (Hatch et al., 2014; Lee et al., 2016; Pagliuso et al., 2018; Manor et al., 2015). Septins (particularly Sept2 and Sept7) are also known to participate in mitochondrial fission via an unknown mechanism (Pagliuso et al., 2016). The dynamin-related protein Drp1 (and likely Dyn2 as well) is predicted to form ring-like or helical oligomers around the fission site to drive membrane constriction (Francy et al., 2015; Ji et al., 2015; Kalia et al., 2018; Mears et al., 2011). The predicted initial diameter of these ring/helical structures is ~100 nm, insufficient to initiate fission for typical mitochondria (>400 nm), so these proteins have been considered late-acting factors. It is likely that Dyn2 drives the final step of scission because it is recruited to mitochondria during the final stages of fission (Lee et al., 2016) and its close homolog, Dyn1, can induce liposome constriction and fission *in vitro* (Sweitzer and Hinshaw, 1998), but this role of Dyn2 is debated (Fonseca et al., 2019). Together, these studies suggest that mitochondrial fission is a stepwise process in which multiple factors act at different stages of constriction. However, loss of any one of these factors does not cause a complete abrogation of fission, suggestive of significant overlaps in their function and therefore not in accordance with a strictly sequential process.

In order to visualize the molecular actors involved in mitochondrial fission directly and artifact-free, we used *in situ* cryogenic <u>e</u>lectron <u>t</u>omography (cryo-ET), a technique that preserves membranes and proteins within cells in near-native fully-hydrated conditions while revealing their 3-dimensional (3-D) architecture at nanometer-scale resolution. Our workflow revealed synchronous constriction of the IMM and OMM, as well as the architecture of ER and cytoskeletal filaments at mitochondrial fission sites, providing mechanistic insights into this highly dynamic and complex process.

## Results

### Capturing mitochondrial fission events by cryo-electron tomography

Cryo-electron tomography (cryo-ET) is a low-throughput technique, and the field of view captured in a 3-D tomogram encompasses only a small fraction of the total cellular volume. Since the population of mitochondria undergoing fission at any point in time is low, randomly imaging these events is impractical. To increase our chances of observing mitochondria undergoing fission, we treated cells (either mouse embryonic fibroblast (MEF) or human bone osteosarcoma epithelial (U2OS) cells) with carbonylcyanide phenylhydrazone (FCCP), a protonophore that causes loss of mitochondrial membrane potential and mitochondrial fragmentation via amplified fission (Cereghetti et al., 2008). To determine the optimal duration of FCCP treatment, we grew MEF cells on glass, treated with FCCP and followed mitochondrial morphology by live-cell fluorescence microscopy for several minutes (Supplementary Fig. 1). By 3-6 minutes post-treatment, very few cells showed hyper-fused mitochondria and there was a concomitant increase in cells displaying fragmented mitochondria (Supplementary Fig. 1A). By 8-10 minutes, a majority of the cells showed fragmented mitochondria, suggesting completion of a large number of fission events. We therefore reasoned that our chances of capturing mitochondria in the act of fission were highest in the 3-6 minute window. This timing is consistent with a previous study showing maximal F-actin association with mitochondria in a similar time frame after FCCP treatment (Li et al., 2015), a result we also observed by staining with FITC-Phalloidin (Supplementary Fig. 1B).

Another major obstacle to cryo-ET imaging is sample thickness. Volumes thicker than ~1 μm are electron-opaque, and ideal samples are < 200 nm thick. In adherent cells, this corresponds to only the thinnest peripheral regions, a small fraction of the total volume. As a result, most of the mitochondrial population is off-limits to direct imaging by cryo-ET. Moreover, FCCP treatment induces a shift in mitochondria localization from the periphery to the perinuclear region (Supplementary Fig. 1B). To target this perinuclear region, we therefore used cryogenic focused ion beam (cryo-FIB) milling to create ~100-300 nm-thick electron-transparent lamellae in the middle of vitrified cells. Our overall workflow is summarized in Fig. 1.

**Figure 1.**
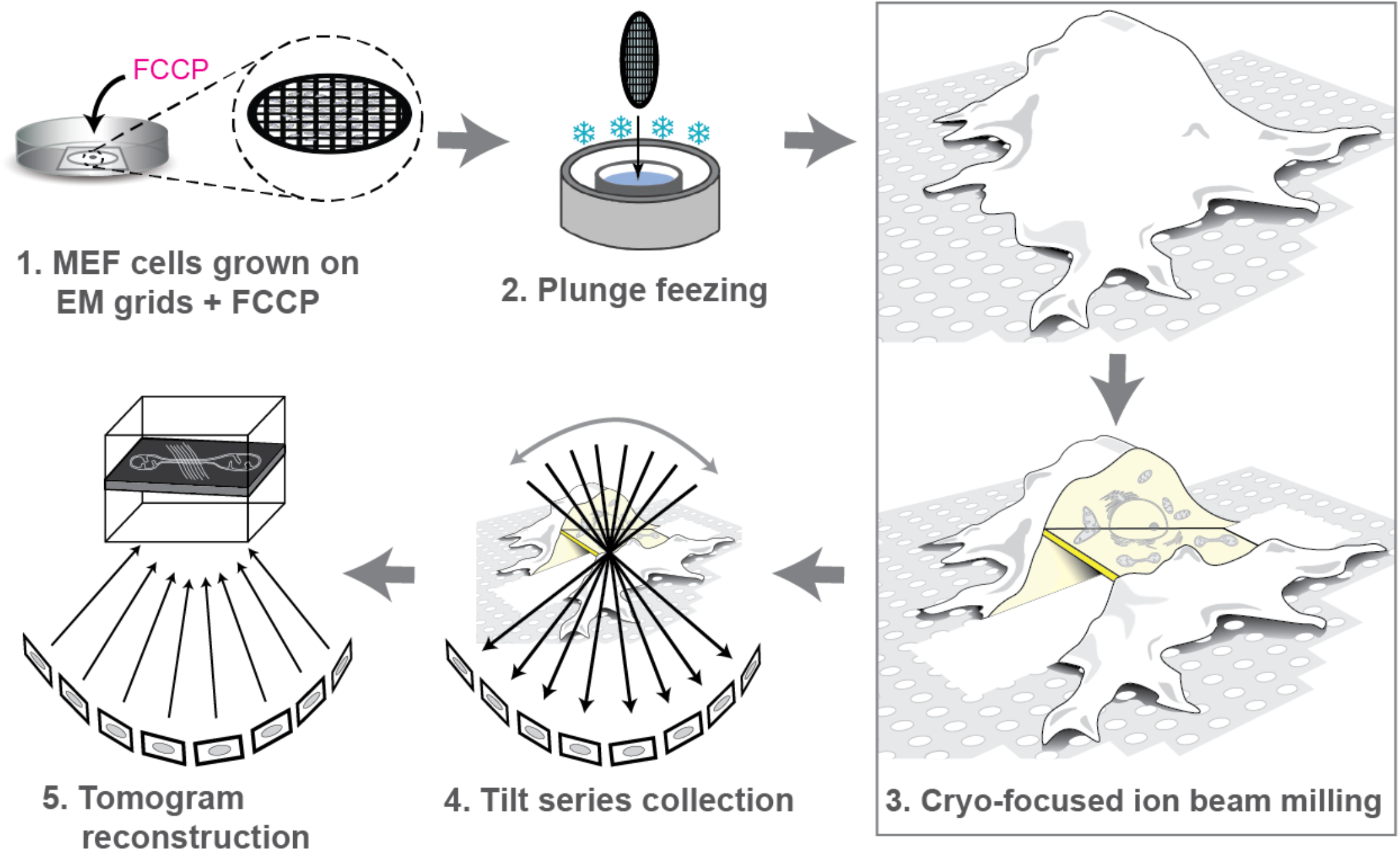
Experimental workflow imaging mitochondrial fission by cryo-ET. **(1)** Mammalian cells were grown on fibronectin-coated EM grids for 24-48 hours and treated with FCCP for 3-6 minutes to induce mitochondrial fission. **(2)** Samples were then plunge-frozen and **(3)** cryo-FIB milled to create thin lamellae randomly positioned near the middle of the cell. **(4)** Lamellae were imaged by cryo-ET to create tilt-series, sets of projection images obtained by iteratively tilting the sample that are used to **(5)** computationally reconstruct a 3-D volume (a tomogram). Images in panel (3) are adapted from (Villa et al., 2013).

Using this workflow to create more than 100 lamellae, we captured mitochondrial constriction events in 18 tomograms of MEF cells and 9 tomograms of U2OS cells. In the more extensive dataset of FCCP-treated MEF cells, the 22 constriction sites (four mitochondria contained two sites) ranged from early stages (light constriction) through late stages (nearly complete constriction). We also identified 3 and 2 constriction events in tomograms of peripheral regions of untreated MEF and INS-1E cells, respectively.

### Conserved ultrastructure of mitochondrial constriction sites

Constrictions in both FCCP-treated and untreated cells of various types (MEF, U2OS and INS-1E) exhibited conserved features (Figs. 2 and 3 and Supplementary Fig. 2) including three different kinds of cytoskeletal filaments, and ER. While the yield from untreated MEF and INS-1E cells was too low for detailed analyses, the overall similarity of the ultrastructure in all cases suggests that our observations in FCCP-treated MEFs are broadly relevant for physiological contexts of mitochondrial fission in mammalian cells.

**Figure 2.**
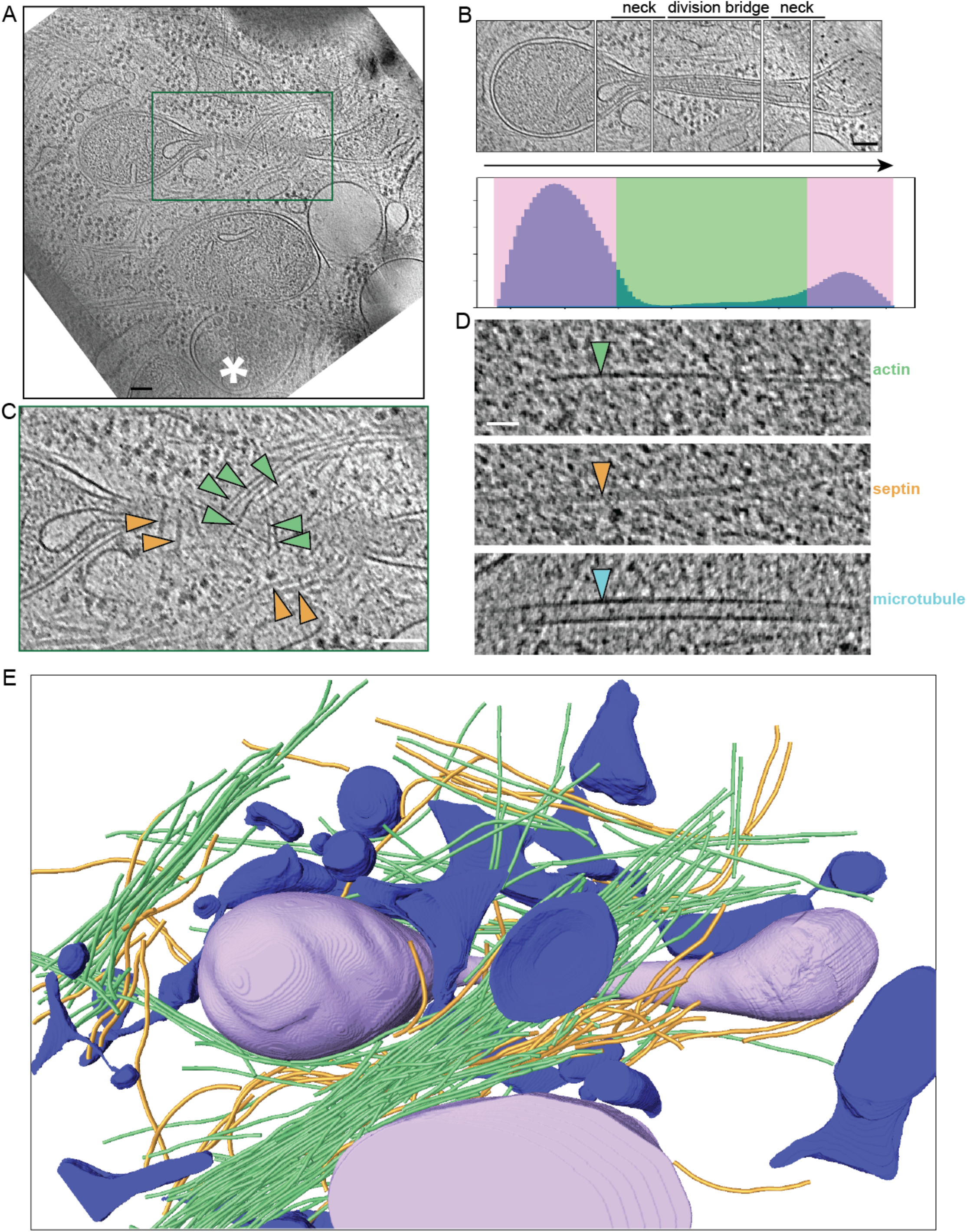
Ultrastructure of mitochondrial constriction sites. (**A**) Slice through a 3-D tomogram of a dividing mitochondrion in an FCCP-treated MEF cell. The asterisk indicates an autolysosome-like organelle. (**B**) Montage of slices at different heights from the tomogram in (A) capturing the central axis of the dividing mitochondrion. The corresponding constriction profile is shown below, depicting the number of voxels contained inside the outer mitochondrial membrane (OMM) in each of 100 sections along its principal axis. This profile allows us to visually classify constriction (green) and non-constriction (pink) zones. (**C**) Enlargement of the boxed region in (A) highlighting filaments visible at the constriction site. (**D**) Cytoskeletal filaments distinguishable in our tomograms: actin (green arrowhead; 7-10 nm diameter); septin (orange arrowhead; 10 nm diameter); and microtubules (teal arrowhead; 24 nm diameter). (**E**) 3-D segmented model of the tomogram in (A) showing OMM (purple), endoplasmic reticulum (ER; blue), actin filaments (green), septin filaments (orange), and microtubule (teal). Scale bars are 100 nm except in (D), which is 20 nm.

**Figure 3.**
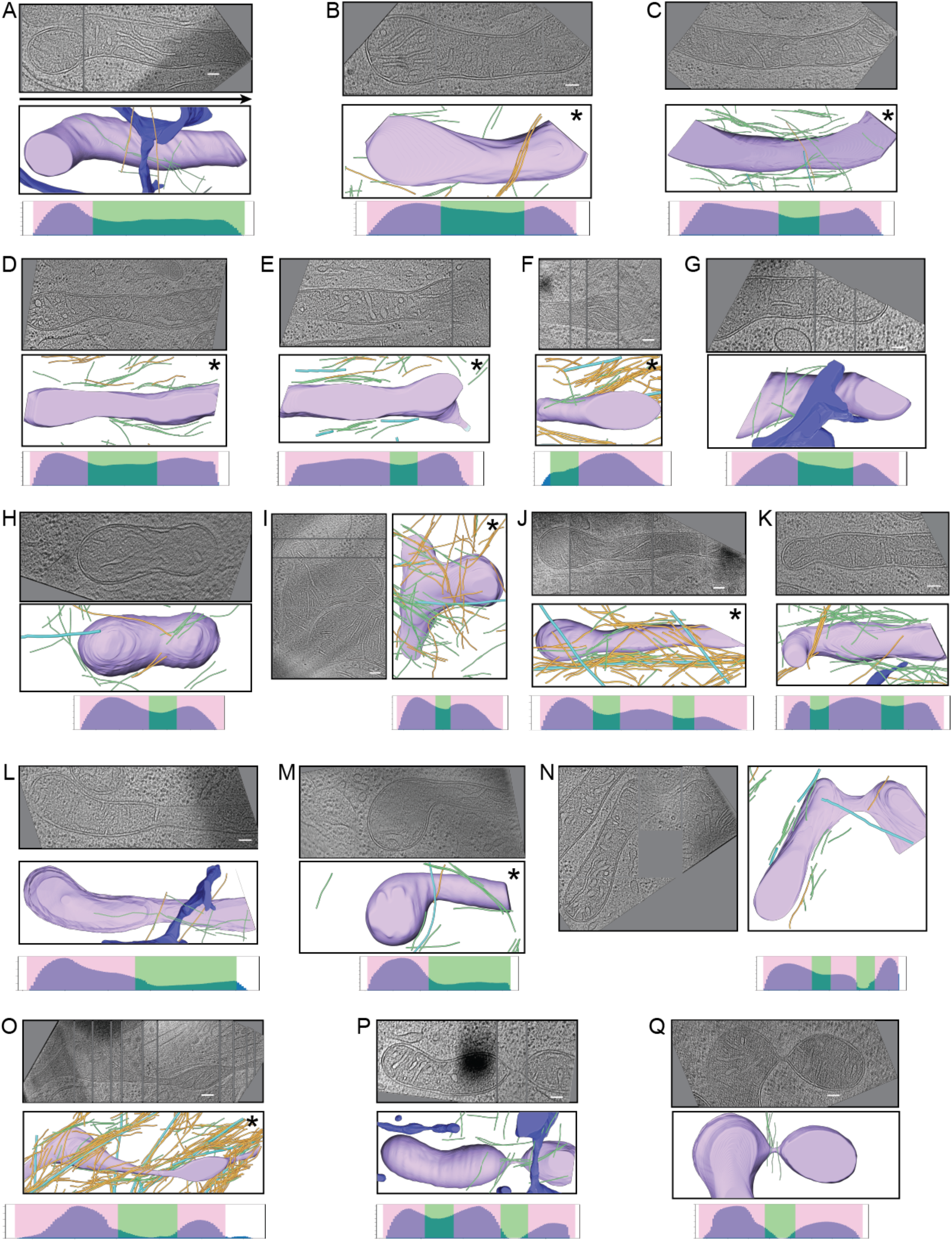
Gallery of mitochondrial constrictions. (**A-Q**) Examples of constricting mitochondria observed in FCCP-treated MEF cells, arranged approximately in order of degree of constriction. In each case, a central tomographic slice (or a montage of multiple slices to capture the central axis) is shown, along with a 3-D segmentation and constriction profile. Color scheme is the same as in Figure 2. Asterisks indicate that ER was not segmented. Scale bars are 100 nm.

Consistent with a previous study (Lee et al., 2016), we observed both focused constriction points and more elongated division bridges (e.g. Fig. 2A, B). Arranging the 18 constricting mitochondria we observed in FCCP-treated MEF cells roughly according to degree of constriction (Fig. 3), we did not observe a correlation between constriction progression and bridge length (visually assessed and highlighted in green in constriction profiles).

We frequently observed ER compartments, sheets and tubes, either smooth ER or decorated with ribosomes, nearby mitochondrial constriction sites. They were derived from more extensive ER tens or hundreds of nanometers away from the constriction sites. In a few cases, we observed close association of ER with a mitochondrial constriction (e.g. Fig. 2E), but we did not observe ER completely encircling any constrictions, contrary to what has been proposed (Hatch et al., 2014; Manor et al., 2015; Phillips and Voeltz, 2016; Rowland and Voeltz, 2012). We also frequently observed what appeared to be autolysosomes in the vicinity of mitochondrial constriction sites (marked with an asterisk in Fig. 2A). These may be involved in the mitophagy that has been observed to follow FCCP-induced fission events (Vives-Bauze et al., 2010).

In addition to these organelles, we observed extensive arrangements of cytoskeletal filaments associated with constriction sites (Fig. 2C). These filaments were of three distinct types (Fig. 2D). Actin filaments were thin and, due to their helical nature, exhibited variable diameter in tomogram slices, between 7 and 10 nm. Microtubules had a uniform diameter of 24 nm. The third class exhibited an intermediate thickness of 10 nm. Due to their known role in mitochondrial fission (Pagliuso et al., 2016), we thought it likely that these were septin filaments. Notably, we did not observe extended helical assemblies consistent with those proposed for dynamins.

### Septins contribute to mitochondrial homeostasis

Mammalian septins – which fall into four families, Sept2, Sept3, Sept6 and Sept7 – assemble into linear apolar filaments composed of a repeating octameric unit (Bertin et al., 2008), which further associate with one another to form higher-order assemblies (Booth et al., 2016; Garcia et al., 2011). We hypothesized that Sept2, a septin known to localize to constriction sites (Pagliuso et al., 2016), is important for the formation of the 10 nm-wide filaments observed in our tomograms. Knocking down Sept2 protein expression in MEF cells (Fig. 4A) caused a dramatic decrease in the abundance of 10-nm filaments in tomograms (Fig. 4B), confirming that these filaments are indeed septins.

**Figure 4.**
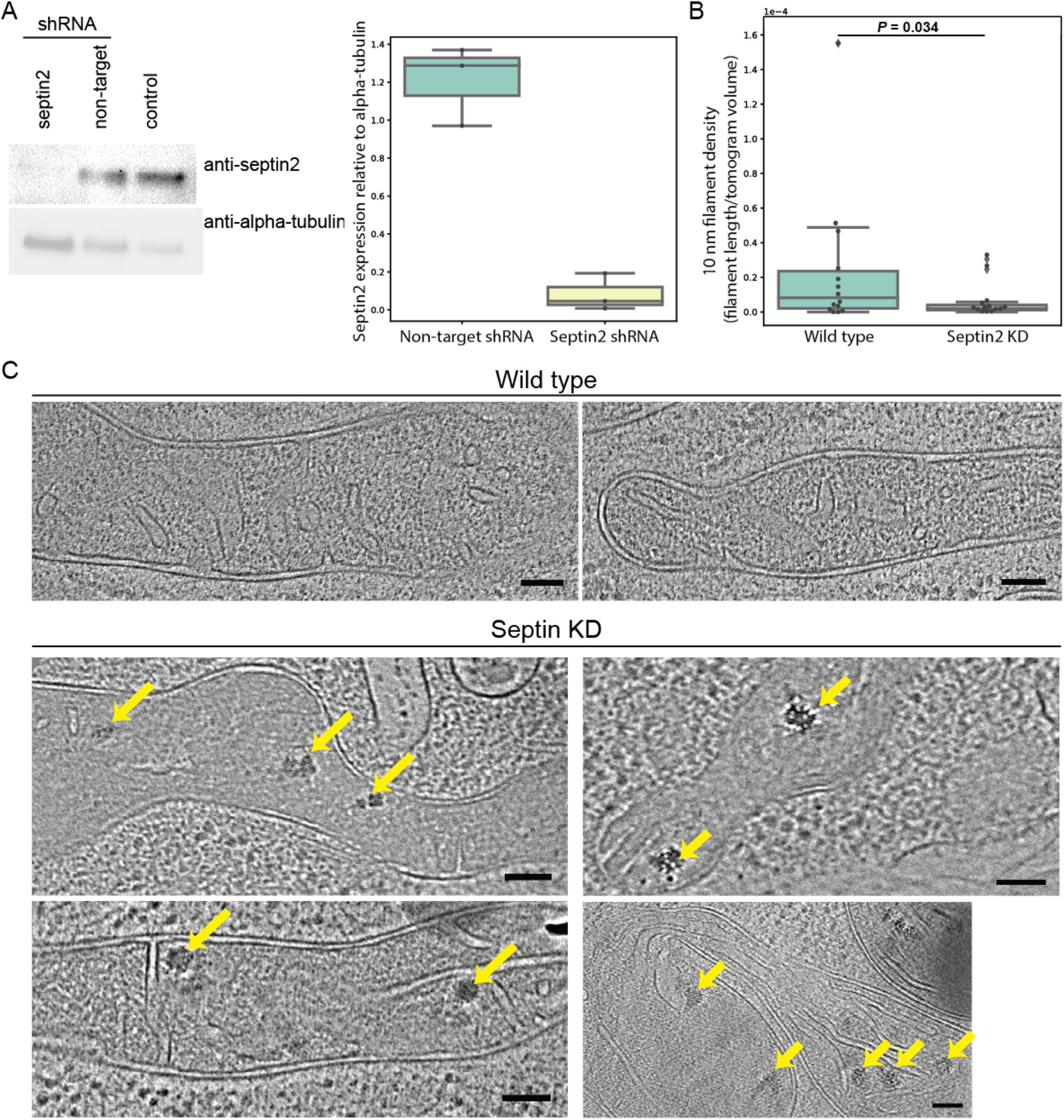
Septin2 knockdown causes loss of 10 nm-wide filaments and alters mitochondrial morphology. (**A**) A representative Western blot showing reduced protein levels of Septin2 in MEF cells following shRNA-mediated knockdown compared to controls (non-targeting shRNA or no shRNA). Alpha-tubulin was used as a loading control. Quantification of three replicates is shown in the box-and-whisker plot at right. In these and subsequent box plots, the central line indicates the median, the box limits the first and third quartile, and the whiskers the minimum and maximum of the dataset, with outliers indicated by diamonds. (**B**) Abundance of 10 nm-wide filaments in tomograms of wild-type and Septin2-knockdown cells following FCCP treatment (n = 14 for each). (**C**) Slices of tomograms showing representative mitochondrial morphology in wild-type (top) and Septin2 knockdown (bottom) cells following FCCP treatment. Note the smoother matrix texture and increased incidence of calcium phosphate granules (yellow arrows). No calcium phosphate granules were observed in any of the 14 wild-type cells. Scale bars are 100 nm.

Interestingly, Sept2 knockdown also altered mitochondrial morphology. A previous EM study identified the form of calcium reserves in mitochondria of MEF cells: solid-state calcium phosphate granules (Wolf et al., 2017). While we did not observe any calcium phosphate granules in FCCP-treated wild-type cells, we observed abundant granules in Sept2-knockdown cells (indicated by yellow arrows in Fig. 4C). In addition, we observed that in these cells, the mitochondrial matrix lost its granular appearance and instead appeared smoothly textured (Fig. 4C). These results suggest that in addition to their role in fission, septins have additional important, but currently unknown, functions in mitochondrial homeostasis.

### Bundles of long unbranched actin filaments mediate mitochondrial fission

F-actin is known to play a key role in mitochondrial fission, although its relevant architecture remains unclear. Consistent with this role, in all tomograms of dividing mitochondria, we observed abundant actin filaments near constriction sites (Figs. 2 and 3, Supplementary Fig. 2). To quantify this association, we demarcated “constriction zones” extending 50 nm out from the OMM at constriction sites. Supporting a specific role in fission, we observed significant enrichment of actin filaments in constriction zones compared to neighboring regions around the mitochondria (Fig. 5A). By contrast, there was no similar enrichment of microtubules (Fig. 5A). Interestingly, we rarely found any free actin filament ends in constriction zones, or even beyond in the larger volume of the tomograms, suggesting that these long filaments are nucleated elsewhere in the cell. Previously, Arp2/3, a major nucleator of actin branching, was shown to be involved in mitochondrial fission triggered by CCCP, a drug similar to FCCP (Fung et al., 2019). The actin filaments we observed, however, were linear with no observable branching, suggesting that Arp2/3 is not organizing F-actin networks locally at division sites.

**Figure 5.**
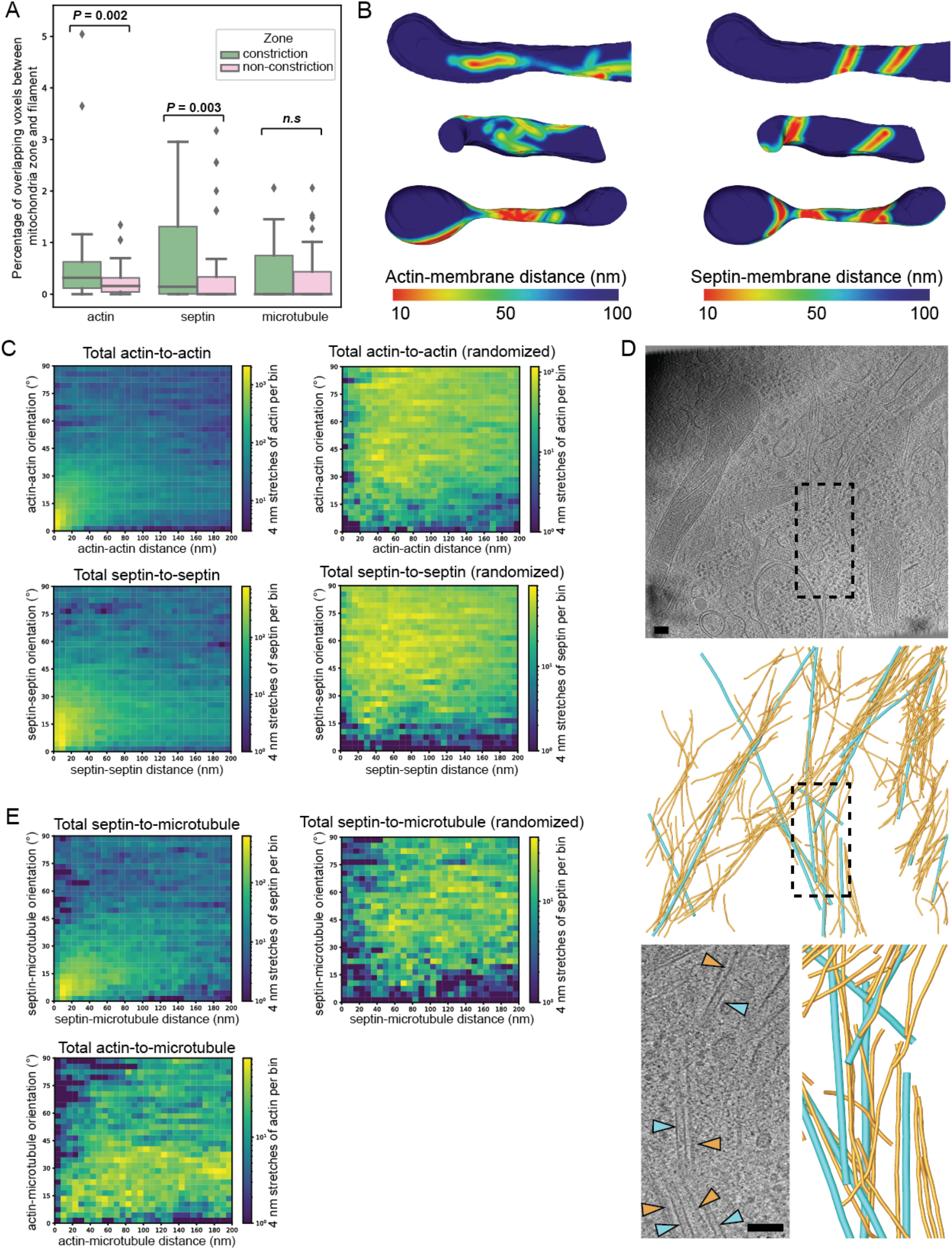
Cytoskeletal filament architecture at mitochondrial constrictions. (**A**) Quantification of filament distribution in constriction and non-constriction zones from 22 mitochondrial constriction sites, showing significant enrichment of actin and septin filaments, but not microtubules. (**B**) 3-D segmented models of the outer mitochondrial membrane (OMM) of the mitochondria shown in Fig. 3M (top), Fig 3G (middle), and Fig. 2 (bottom). Surfaces are colored according to the proximity of actin (left) and septin (right) filaments. (**C**) 2-D histograms of the relative orientations and distances between neighboring segments of actin (left panels; from 13 tomograms) and septin (right panels; from 11 tomograms), showing a strong correlation compared to simulated negative controls with randomized positions and orientations. (**D**) Slice from a tomogram and 3-D segmented model showing close parallel association between septin filaments (orange) and microtubules (teal). The boxed region is enlarged on the right. (**E**) 2-D histograms of the relative orientations and distances between microtubules and septin filaments (left panels; from 10 tomograms) or actin filaments (right panel; from 11 tomograms). Septin, but not actin, filaments show a strong positional and orientational correlation compared to the randomized negative control. Scale bars are 100 nm.

Actin filaments were oriented mostly obliquely or perpendicular to the long axis of the dividing mitochondria, and were closely associated with the OMM, often within 10 nm (Fig. 5B). They also displayed a tendency to bundle, a phenomenon we quantified by measuring the distances and relative orientations in 3-D between short (4 nm) segments of neighboring actin filaments. The resulting heatmaps revealed a tight correlation in orientation of adjacent filaments, consistent with bundling (Fig. 5C). As a negative control, we simulated random positions and orientations for the same actin filaments, which caused the relationship to disappear (Fig. 5C).

### Septin bundles are found at mitochondrial necks

While septins have been implicated in mitochondrial fission, the mechanism is unknown. As described above, we observed numerous long, unbranched septin filaments in our tomograms of constriction sites (Figs. 2 and 3, Supplementary Fig. 2). They were present from the earliest stages of constriction and persisted throughout the process, missing only from the final stage, in which the IMM had separated and only a bridge of OMM remained connected (Fig. 3P and Q). Similarly to F-actin, no septin filament ends were detected, suggesting they were long filaments that originated far from the mitochondria, were arranged obliquely or perpendicular to the long axis of the mitochondria, were significantly enriched in constriction zones (Fig. 5A), and closely associated (≤10 nm) with the OMM (Fig. 5B). Also similarly to actin, septin filaments formed tight bundles of parallel filaments (Fig. 5C). Septin bundles often partially tracked the local curvature of the OMM (e.g. Fig 3K). However, they did not wrap to encircle constrictions. In mitochondria with long division bridges, we observed preferential localization of septin bundles to the “neck” regions on either side (or, if both were not visible in the tomogram, at least one side) of the bridge (Fig. 5B). This localization pattern was notably distinct from that of F-actin (Fig. 5B).

### Septins interact with microtubules at constriction sites

Intriguingly, we noticed septin bundles interacting with microtubules, both inside and outside constriction zones (e.g. Fig. 3F, J and O). When we mapped all septin filaments and microtubules in our tomograms, they exhibited a strong tendency to interact laterally (Fig. 5D). Quantifying the relationship between distance and orientation of segments of adjacent septin filaments and microtubules, we observed a strong correlation in orientation (Fig. 5E). No similar relationship was observed between actin filaments and microtubules in our tomograms (Fig. 5E). Since septins, but not microtubules, localized preferentially to constriction zones (Fig. 5A), this suggests that they may be somehow guiding microtubules in the vicinity to fission sites.

### Inner and outer membrane constriction is synchronized throughout mitochondrial fission

While, of course, IMM separation must precede that of the OMM, the extent to which constriction of the two membranes is coordinated during fission is unknown. It has been proposed that constriction of the IMM occurs independently of OMM constriction, via a dedicated mechanism (Chakrabarti et al., 2018; Cho et al., 2017). Our tomograms resolved both membranes well and revealed a high degree of synchrony throughout constriction (Figs. 3 and 6A). This coupling is consistent with constrictive force being applied from outside the OMM, transmitted indirectly to the IMM through bridging proteins. We observed coordinated constriction until the IMM was brought into contact, pinching off and sealing the new compartments on each side (Fig. 6B). At this point, the remaining neck of OMM was ~25 nm wide (Fig. 6B).

**Figure 6.**
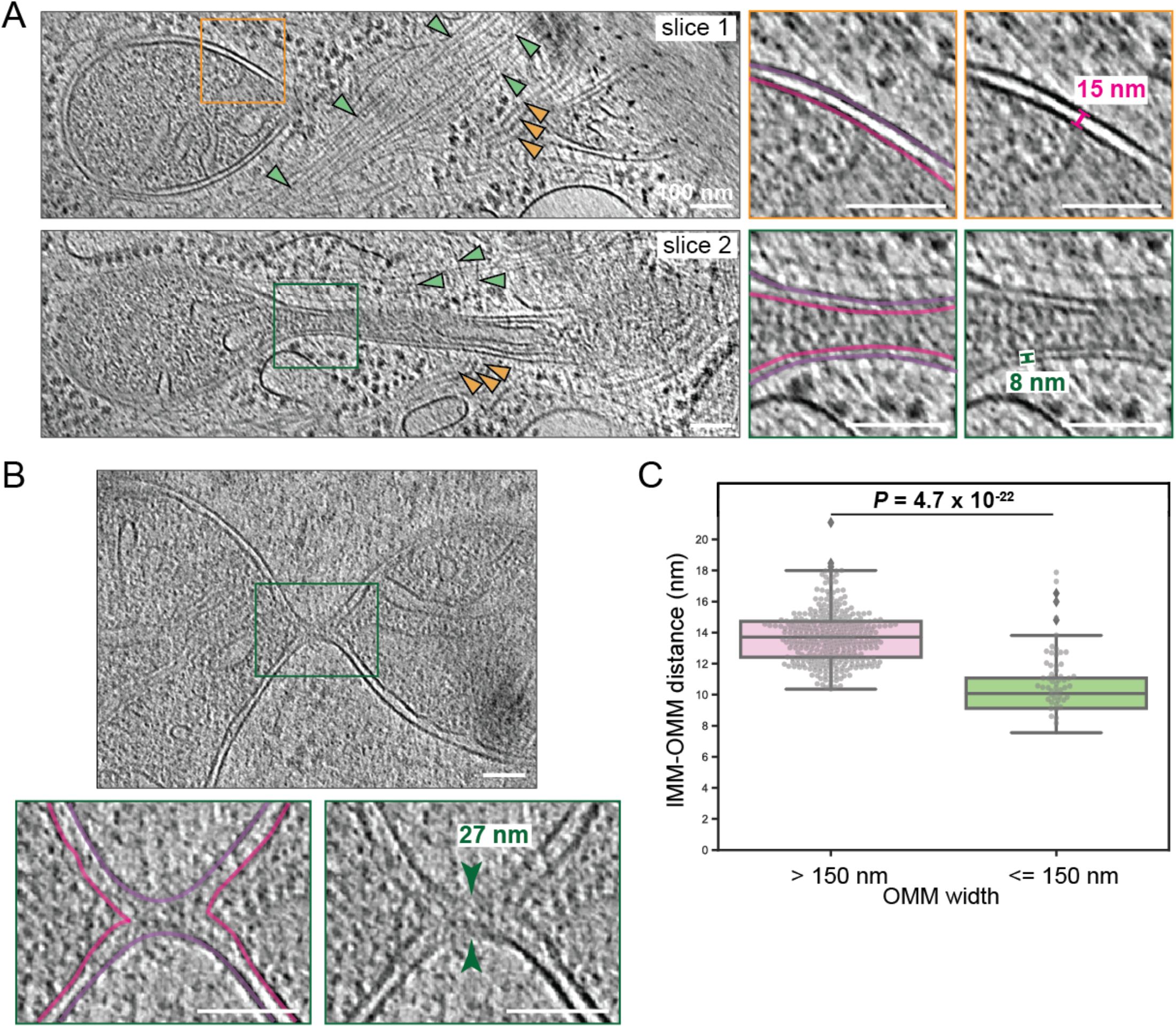
Synchronous constriction of mitochondrial membranes. (**A**) Tomogram slices at different heights through the constricting mitochondrion shown in Fig. 2. Actin and septin filaments are indicated by green and orange arrowheads, respectively. Enlargements of the boxed regions are shown at right, highlighting the spacing between the inner mitochondrial membrane (IMM; pink) and outer mitochondrial membrane (OMM; purple). (**B**) Tomogram slice of the constricting mitochondrion shown in Fig. 3Q. An enlargement of the boxed region is shown below, highlighting complete fission of the IMM (pink) but not OMM (purple). (**C**) Box-and-whisker plot of measurements of IMM-OMM distances at multiple positions along 15 mitochondrial constriction profiles with a total width (measured from the center of the OMM on either side) of either >150 nm (pink) or ≤150 nm (green). Scale bars are 100 nm.

Interestingly, we observed a change in the distance between the IMM and OMM at later stages of constriction (Fig. 6A). We quantified this effect by measuring the intermembrane distance at multiple points along 15 constriction sites. We found that in early (light) constrictions, the intermembrane distance remained unchanged (~14 nm). However, in deeper constriction profiles with a width of 150 nm or less, this distance decreased to ~10 nm (Fig. 6C).

## Discussion

Mitochondria retain many characteristics of bacteria, reflecting their endosymbiotic origin. Mitochondrial fission is therefore often compared to bacterial cell division, which is driven by machinery from within exerting a pulling force on the membrane. Mitochondria, however, lack a homologue of the force-generating protein FtsZ, making it unlikely that a similar mechanism exerts a constrictive force on the IMM. Instead, force may come either from constriction of the OMM or through an independent dedicated mechanism, as has been proposed recently (Chakrabarti et al., 2018; Cho et al., 2017). Our observations here of synchronous membrane constriction and compression of the intermembrane space at the later stages of fission are consistent with a model in which IMM constriction is driven predominantly by OMM constriction via flexible protein spacers between the two membranes such as the mitochondrial intermembrane space bridging complex (Ott et al., 2012). This also suggests a change in forces or kinetics at the later stages, perhaps associated with recruitment of known late-acting factors such as dynamin. When the channel inside the IMM becomes sufficiently narrow (on the order of a few nanometers), the IMM fuses, presumably followed by a similar process in the OMM to complete fission.

How is constrictive force applied to the OMM? In our tomograms, cytosolic factors previously implicated in mitochondrial fission – ER, actin and septin – exhibited no clear association with particular stages of constriction, contrary to a sequential model of factor recruitment (Hatch et al., 2014). Previous studies using fluorescence microscopy and biochemical assays found that ER-mitochondria contact sites coincide with sites of mitochondrial fission and mtDNA replication (Friedman et al., 2011; Lewis et al., 2016), implicating ER in mitochondrial constriction, either with or without a contribution from actin filaments. In the latter model, ER tubules are proposed to wrap around the mitochondria, initiating constriction on their own (Rowland and Voeltz, 2012). However, our observations of only partial ER coverage around mitochondrial constrictions make this model for force generation unlikely, or at least context-specific. In the alternative model, ER-mitochondria contact sites generate constrictive force by stimulating local actin polymerization. Consistent with this model, an ER-specific isoform of INF2, a formin protein that nucleates actin filaments, has been shown to be important for mitochondrial fission (Korobova et al., 2013). In addition, Spire1C, a formin-domain-containing protein, has been shown to localize to the OMM surface, where it interacts with INF2 on the ER and enhances actin nucleation, thereby promoting ER-mediated mitochondrial fission (Manor et al., 2015). Thus actin filaments are strongly implicated in the fission process.

At least two models, which are not mutually exclusive, have been proposed to explain how actin filaments could generate the force required for constriction. In one model, actin filaments nucleated at the ER polymerize towards the mitochondrion, applying an inward force to points on the OMM surface (Manor et al., 2015). In the other model, contact-site ER nucleates actin filaments around the mitochondrion. Bipolar myosin II molecules then bind filaments from different nucleation points, walking them together to constrict the ER, and applying force to the trapped OMM (Hatch et al., 2014). However, a recent platinum replica electron microscopy (PREM) study revealed an interstitial network of actin filaments at mitochondrial constriction sites, along with a stochastic distribution of myosin II (Yang and Svitkina, 2019). This arrangement is inconsistent with either of the two models just described, and the authors instead proposed that bipolar myosin activity might produce heterogeneous deformations in the actin network that exert local force on trapped OMM. In that study, however, samples were prepared by treatment with saponin (a mild detergent) and mechanical unroofing to gain access to the cell interior, so it is unclear how well the observations reflect the structure of unperturbed cells.

Our use of cryo-FIB milling and cryo-ET, which better preserve cellular membranes and proteins, gave us a window into the native arrangement of actin at mitochondrial constriction sites. We observed actin filaments neither encircling fission sites nor originating from ER at contact sites, but instead most consistent with the network arrangement proposed in the PREM study (Yang and Svitkina, 2019). The enhanced resolution of our tomograms also enabled additional observations about actin filaments: (i) they displayed abundant bundling; (ii) they were significantly enriched close (≤10 nm) to the OMM at constriction sites; (iii) they were unbranched and long (not originating at nearby ER membranes); and (iv) their nucleation (by formins or Arp2/3) did not seem to be locally controlled at constriction sites. Unfortunately, we were unable to identify myosin II molecules in our tomograms; the slender molecules are difficult to resolve, with their small heads tightly interacting with actin filaments and only a coiled-coil linker region in between. While bipolar myosin may be mediating actin-driven constriction, it is also possible that the actin filaments are acting as dynamic reservoirs for recruiting dynamin (or dynamin-like) proteins to the OMM. Actin filaments have previously been shown to bind Drp1 and enhance its oligomerization at constriction sites (Ji et al., 2015), and Drp1 has also been shown to increase actin filament bundling (Hatch et al., 2016; Ji et al., 2015), which we observe consistently in our tomograms.

It is noteworthy that we could not identify dynamins at constriction sites, even in late stages when we would expect Drp1, and possibly Dyn2 (its role in fission is debated), to be present (Fonseca et al., 2019; Lee et al., 2016). While it is known that dynamins form oligomers to drive constriction and scission, the exact nature of the assemblies remains unclear. Like Dyn1, Dyn2 has been hypothesized to form helices around mitochondrial constrictions (Chappie et al., 2011; Pucadyil, 2011; Ramachandran and Schmid, 2018); Drp1 has been proposed to form either similar helices or rings (Ji et al., 2015; Kalia et al., 2018; Mears et al., 2011). However, our tomograms showed no repetitive helical structures at mitochondrial constrictions, suggesting that neither protein forms an elaborate, stable helical assembly *in vivo*.

Septins were recently discovered to participate in mitochondrial fission (Pagliuso et al., 2016), but the details of their involvement, including any interplay with actin, remain unknown. In our tomograms, we observed bundles of long, unbranched, Sept2-dependent filaments, concentrated at constriction sites. Their arrangement, frequently localized to the neck region of division bridges, differed from that of actin filaments, suggesting distinct roles. Septins are known to sense micron-scale membrane curvature (Bridges et al., 2016; McMurray, 2019), including at the poles of rod-shaped bacteria (Krokowski et al., 2018). It is therefore possible that the septin filaments we observed in our tomograms are sensing membrane curvature, if not driving remodeling directly, then perhaps reinforcing it by physically preventing the OMM from relaxing back to a larger diameter. Consistent with such a role, septins are known to participate in another membrane deformation process: cytokinesis in yeast (Bertin et al., 2012; Ong et al., 2014). In that system, septins organize into diverse assemblies including long linear filaments, circumferential rings, and a gauze-like meshwork. The functional implications of these arrangements are unclear, however, and our observation of linear septin filaments involved in mitochondrial fission may shed light on their role in other membrane remodeling processes as well. In addition to potentially directly driving membrane constriction, septins may also recruit Drp1 to constriction sites to finish the process (Pagliuso et al., 2016).

Interestingly, our tomograms revealed interactions between septin filaments and microtubules at mitochondrial constriction sites. It is known that septins can spatially guide microtubules and their plus-end dynamics in epithelial cells (Bowen et al., 2011), and this function could be more universal. In the case of damaged mitochondria undergoing mitophagy, septins could direct microtubules to fission sites to facilitate delivery to the autophagy pathway. A previous study found that damaged mitochondria are transported in pieces away from sites of mitochondrial fission (Yang and Yang, 2013), and in our tomograms we observed autolysosomes near fission sites. In the future, it will be interesting to apply the workflow we developed here for nanoscale *in situ* cryo-ET imaging to downstream processes following mitochondrial fission such as this targeted mitophagy.

## Materials and Methods

### Cell growth

Adherent mouse embryonic fibroblasts (MEF; gifts of D. Chan, California Institute of Technology and G. Shadel of Salk Institute of Biological Studies), bone osteosarcoma cells (U2OS; gift of R.L Wiseman, Scripps Research Institute), and rat insulinoma cells (INS-1E; gift of P. Maechler, Université de Genève) were grown in a humidified 37 °C incubator with a constant supply of 5% CO_2_. Cells were cultured in high glucose (L-Glutamine +) Dulbecco’s modified Eagle’s medium (DMEM; DML09, Caisson Labs, Smithfield, UT) supplemented with 10% fetal bovine serum (FBS; Cat No – 10437028, ThermoFisher), 1 mM sodium pyruvate (Cat No – 11360070, ThermoFisher), 100 units/mL penicillin and 100 μg/mL streptomycin. Plasmids for shRNA-mediated knockdown of Sept2 and Dyn2 were maintained in MEF cells using 3 μg/mL puromycin (gift of D. Chan). For cryo-FIB milling and cryo-ET, cells were grown on 200 mesh gold R2/2 London Finder Quantifoil grids (Quantifoil Micro Tools GmbH, Jena, Germany). Prior to seeding cells, these grids were coated with 0.1 mg/mL human fibronectin (Cat No – C-43060, PromoCell) by floating them on fibronectin droplets on parafilm for approx. 15-30 min. Cells were grown to a density of approximately two to three per grid-square over a period of one to two days.

### Gene knockdown

A knockdown cell line for Sept2 was created in MEF cells using retroviral transduction with guidance from C.-S. Shin and reagents kindly gifted by D. Chan, both at California Institute of Technology. Briefly, an shRNA fragment for Sept2 (a hairpin forming sequence derived from the Sept2 targeting sequence of 5’ GCAGTTTGAACGCTACCTACA 3’) was inserted into pRetroX-H1 and confirmed by sequencing. For delivery of this shRNA plasmid into MEF cells, retroviruses (packaged with this plasmid) were produced by co-transfecting 293T cells with the retrovirus packaging vector, pCL-Eco^2.1^, and the shRNA plasmid. Viruses were harvested from supernatant by centrifugation (to remove cell debris) and filtration through a 0.45 μm syringe filter. MEF cells were subsequently infected with these retroviruses at a low confluence (~10%) in the presence of 4 μg/mL polybrene to enhance virus-host interaction; after the addition of viruses to cells, low speed centrifugation was performed to increase infection efficiency. MEF cells successfully transduced with the shRNA plasmid were selected using 3 μg/ mL puromycin and maintained in the presence of this drug. Sept2 knockdown was confirmed by western blotting using a rabbit monoclonal primary antibody against Sept2 [EPR12123] (Cat No – ab179436, Abcam, Cambridge, MA; used at 1:500-1:2000 dilution) and a goat anti-rabbit IgG (H+L) secondary antibody conjugated with horseradish peroxidase (HRP) (Peroxidase AffiniPure; Cat. No – 111-035-003, Jackson ImmunoResearch, PA, USA; used at 1:10000 dilution). A mouse anti-alpha-tubulin primary antibody (monoclonal; Cat No – T6199, Sigma-Aldrich, MO, USA) and a goat anti-mouse secondary antibody conjugated with HRP (Cat. No – 115-056-003, Jackson ImmunoResearch) were used to quantify α-tubulin in the same samples as loading control. The band intensities representing Sept2 protein expression level were quantified and normalized to the band intensities representing alpha-tubulin. The intensities of protein bands from 3 different experiments were quantified using the ImageJ Gel Analysis program as illustrated here (https://imagej.nih.gov/ij/docs/menus/analyze.html#gels and https://lukemiller.org/index.php/2010/11/analyzing-gels-and-western-blots-with-image-j/). As shown in Fig. 4A, there is significantly less Sept2 expression relative to alpha-tubulin in shRNA-mediated knockdown. The *p*-value was calculated using the Mann-Whitney rank test, a non-parametric alternative to the independent Student’s t-test.

### Confocal light microscopy

#### Experimentation

In order to capture mitochondrial fission events by cryo-electron tomography (cryo-ET), we first monitored mitochondrial fragmentation using live-cell light microscopy in MEF cells. Imaging was performed at the Caltech Biological Imaging Facility on a Zeiss LSM800 microscope equipped with a large environmental chamber to maintain the temperature at 37°C and a smaller insert module that helped maintain both the temperature and a CO2 level of 5%. MEF cells were first treated with 200 nM of MitoTracker Red CMXRos (Cat No - M7512, ThermoFisher) to fluorescently label mitochondria for confocal light microscopy. Cells were treated in full medium (no HEPES) for 15 min at 37 °C and washed twice using the same medium (no dye) before treatment with 10 μM FCCP. For experiments involving visualization of F-actin, cells were treated with FITC-phalloidin (Cat No - ALX-350-268-MC01, Enzo Life Sciences, Ann Arbor, USA). Briefly, post-FCCP-treatment, cells were fixed with 4% paraformaldehyde (PFA) in phosphate buffer saline (PBS) for 15 min, washed twice with Hank’s Balanced Salt solution (HBSS), incubated in acetone for 4 min at −20 °C, washed twice again with HBSS, stained with 220 nM of FITC-phalloidin for 20 min, and finally washed twice with HBSS. Both bright-field and fluorescence imaging were performed using an LD C-Apochromat 40× water-immersion objective with a numerical aperture of 1.1, and images were recorded using photomultiplier tubes (PMTs; for bright-field image) and GaAsP-PMT (for fluorescence). FITC-phalloidin and MitoTracker Red were imaged using diode lasers at 488 nm and 561 nm, respectively, operating at ~1.5 to 2% of its maximum power. The maximum power for the laser lines was 500 mW at the source but measured to be ~750 μW at the level of the objective lens for the 488-nm laser and ~500 μW for the 561-nm laser. Mitochondrial morphology was monitored ~every 2 min for up to 30 min after FCCP treatment. A scan speed of 7 was used for imaging cells. The pixel size for imaging was set at 0.312 μm (0.156 μm at a zoom factor of 2), while the image sizes were fixed at 512 by 512 pixels.

#### Quantification of mitochondrial morphology

We collected images of cells with fluorescently-labeled mitochondria at various time points post-FCCP treatment. We blinded these images and scored the cells for containing primarily hyperfused, intermediate, or fragmented mitochondrial morphology (see Supplementary Fig. 1A for examples of mitochondrial morphology classifications). A total of 174 cells distributed across different time points were analyzed for this purpose.

### Plunge-freezing

EM grids containing adherent cells were plunge-frozen in a liquid ethane/propane mixture (Tivol et al., 2008) using a Vitrobot Mark IV (FEI, Hillsboro, OR). The Vitrobot was set to 95-100% relative humidity at 37 °C and blotting was done manually from the back side of the grids using Whatman filter paper strips. In the few cases where cryo-ET was performed on peripheral cell regions without cryo-FIB milling (INS-1E and untreated MEF cells), Au fiducials (Cat No – 15703, Ted Pella) resuspended in 1% bovine serum albumin (BSA) were added to the grids just before blotting and plunge freezing. Plunge-frozen grids were subsequently loaded into Krios autogrid cartridges (ThermoFisher). EM cartridges containing frozen grids were stored in liquid nitrogen and maintained at ≤-170 °C throughout storage, transfer and cryo-EM imaging.

### Cryo-focused ion beam milling (cryo-FIB milling)

These experiments were performed at two different facilities - (1) at the Transmission Electron Microscopy Facility for material science at Caltech using a Versa 3D DualBeam scanning electron microscope (ThermoFisher) equipped with a cryo-stage and a cryo-transfer system (PP3010 from Quorum Technologies Ltd., East Sussex, United Kingdom); and (2) the Scripps Research Institute Hazen Cryo-EM Microscopy Suite using an Aquilos 1 cryo-DualBeam FIB/SEM. Plunge-frozen cells were first coated with a 5-15 nm platinum (Pt) layer using a sputter coater. Ideally positioned cells were first identified on the grids (preferably in the middle of the grid squares) using the electron beam (5 kV for observing topology and 20 kV for observing grid bars). Lamellae were milled at an angle of ~15° with respect to the horizontal (corresponding to 22° on the cryo-stage) or less. After tilting the stage to the desired angle, material above and below the target volume was milled away by progressively stepping down the FIB current (at a constant voltage of 30 kV) as we approached the target volume: starting from 300-100 pA for removing material up to ~1.5 μm from the target volume and finishing the process with a polishing step using 10-30 pA. The electron beam was used intermittently to monitor the milling process. Different voltages ranging from 5 kV to 20 kV helped discern the topology of cells and their positioning on the grids. We prepared more than 100 lamellae in this manner for this study.

### Cryo-electron tomography (cryo-ET)

Cryo-ET was performed using a ThermoFisher Krios G3i 300 kV FEG cryo-TEM at the Caltech CryoEM Facility and using a similar instrument at the Scripps Research Institute Hazen Cryo-EM Microscopy Suite. The microscopes were equipped with a 4k x 4k K2 Summit direct detector (Gatan, Inc.) operated in electron counting mode. For data collected at the Caltech CryoEM facility, an energy filter was used to increase the contrast at both medium and higher magnifications with a slit width of 50 eV and 20 eV, respectively. Additionally, defocus values of close to negative 100 and negative 1-to-8 μm were used to boost the contrast (in the lower spatial resolution range) at the medium and higher magnifications respectively. Magnifications typically used on the Krios were 3,600 X (or 4,800 X; in the medium range) and 26,000 X (in the higher range), corresponding to pixel sizes of 4.2 nm/3.1 nm and 5.38 Å respectively. For data collected at the Caltech CryoEM facility, a Volta phase plate (VPP) was optionally used to further improve contrast at higher magnifications in certain cases. SerialEM software (Mastronarde, 2005) was used for all imaging. The order of imaging steps was as follows. First, a full grid montage at a low magnification (close to 100 X) was acquired to locate the milled lamella on a grid. Mitochondria were then identified in the lamella by their characteristic appearance (double membrane and matrix morphology) in projection images. Once the areas of interest were identified and marked, anchor maps were used to revisit these locations and collect tilt-series in an automated fashion. Each tilt-series was collected from negative 60° to positive 60° with an increment of 2° using the low dose functions of tracking and focusing. The cumulative dose of each tilt-series ranged between 80 and 150 e^-^/Å^2^. Once acquired, tilt-series were binned into 1k x 1k arrays before alignment and reconstruction into 3D tomograms with the IMOD software package (Kremer et al., 1996) and tomo3D (Agulleiro et al., 2011). Tilt series were aligned using 10 nm Au fiducials (for unmilled samples) or patch tracking (for milled samples) in IMOD while reconstructions were performed using SIRT in tomo3D.

### Segmentations and quantifications

#### Segmentation of mitochondrial and ER membranes

Preliminary segmentations were performed using IMOD before the final segmentations were performed using AMIRA (ThermoFisher), in both cases, manually. The final segmentations of a total of 22 mitochondrial constriction sites along 18 different mitochondrial profiles are included in this study.

#### Segmentation of actin, septin and microtubule filaments

For all figures (except Supplementary Fig. 2C), actin and septin filaments were segmented using automated detection and tracing using the XTracing for Amira extension package, which implements methods described in (Rigort et al., 2012). Separate cylinder correlation procedures were performed to detect actin filaments (Cylinder length: 500; Angular sampling: 5; Mask cylinder radii: 70; Outer radius: 40; Inner radius: 0) and septin filaments (Cylinder length: 400; Angular sampling: 5; Mask cylinder radii: 100; Outer radius: 60; Inner radius: 0). Next, interactive thresholding was performed on the output of the cylinder correlation to identify ideal thresholds where all parts of filaments are visible. These threshold values were used to determine the parameters for minimum and continuation correlation values for the Trace correlation lines procedure (Direction coefficient: 0.2; Minimum distance: 60; minimum length: 2500; Search cone: 300; Search angle: 30). The results of the tracing procedure were evaluated in Amira and false-positives were removed using the Filament editor. For supplementary Fig. 2C, actin and septin filaments were segmented manually (the signal-to-noise in the tomogram did not allow for exhaustive segmentation of these filaments). Microtubules were traced manually using the manual filament tracer option in the Amira Filament editor window.

#### Quantification of filaments at mitochondrial constriction sites

A region around mitochondria up to 50 nm from the outer mitochondrial membrane (OMM) was analyzed to quantify association of filaments with mitochondria. This region was first classified into constriction and non-constriction zones in a semi-automated fashion as follows. The principal axis of each mitochondrion was computed by principal component analysis on the coordinates of all of its segmented voxels. This principal axis was subsequently divided into 100 sections along its length and mitochondrial volumes constituting each section were plotted against the position of those sections along the principal axis. The resulting plot helped to visualize the constriction profile for each mitochondrion, which in turn helped to manually classify the surrounding region into constriction and non-constriction zones as shown in Figs. 2 and 3. The abundance of a certain type of filament in each of the zones was computed as the percentage occupancy for that filament type in the total number of voxels contained in that zone. After repeating the procedure for 19 mitochondrial profiles, the *p*-value was calculated using the Wilcoxon signed rank test, a non-parametric alternative to the paired Student’s t-test.

#### Quantification of inter-filament distances and orientations

For each segmented tomogram, filaments in each class (microtubule, septin, and actin) were subdivided into 4 nm segments using linear interpolation, and both the position and orientation of these segments were recorded as numpy arrays. For each segment on each filament, the distance was measured to every segment on every other filament of the same class and every filament of a different class. The distance as well as the calculated relative orientation of the nearest overall segment for each class was recorded. These measurements were aggregated for all filaments of each class from all tomograms (n = 13, 11, 10, and 11 tomograms for actin-actin, septin-septin, septin-microtubule, and actin-microtubule interactions, respectively) in order to generate the 2-dimensional histograms in Fig. 5.

As a negative control for filament-filament interactions, the same analyses were run with randomized filament positions and orientations. To do this, after interpolation, all of the filaments were aligned by their first segment and then randomized such that that first segment was oriented ±30° relative to the growth plane (but otherwise unconstrained). The starting orientations of the filaments were then randomized, and in order to maintain constant total filament density, they were wrapped using a repeating boundary condition, with distance measurements only considered within the tomogram rather than across boundaries. Each filament maintained the same shape as prior to randomization, thus enabling randomization while maintaining physiological curvatures.

Interpolation and distance measurement were calculated with optimized functions in scipy {{32015543}} and using numpy arrays {{32939066}} to maximize performance, allowing comparison of hundreds of microns of interacting filaments in minutes on a single core of a consumer laptop CPU. Plotting was performed with matplotlib {{doi:10.1109/MCSE.2007.55}}. The python scripts to run these analyses are available at https://github.com/Grotjahnlab/measure_models and are free to use or modify with a BSD license.

#### Generation of heat maps for outer mitochondrial membrane (OMM) proximities to filaments

Prior to distance calculations, all segmented models were converted to surfaces in Amira using either the “Generate surface” function for voxel-based OMM segmentations or the “Extract surface from spatial graph” function for filament segmentation in Amira. The “surface distance” function in Amira was used to calculate the distance between segmented surface models of the OMM (surface 1) and cytoskeletal filaments (surface 2). For visual representation of the surface proximity heat maps, the OMM was colored by setting the “color field” to the output from the “surface distance” calculation, and the “color map” to “Physics”. The range of the color map was adjusted to reflect appropriate minimum and maximum distances.

#### Quantification of abundance of 10 nm filaments (septin) in wild-type and Sept2 knockdown cells

Automated detection, tracing, and segmentation of these cytoskeletal filaments was performed using the XTracing for Amira extension package using the same parameters for septin filaments (as described above) for both wild-type (n=14 tomograms) and Sept2 knockdown (n=14 tomograms) cells. The total length of these filaments was calculated and divided by the total volume of the tomogram to calculate their total abundance in wild-type and Sept2 knockdown cells. The *p*-value was calculated using the Mann-Whitney rank test.

#### Quantification of intermembrane distance between IMM and OMM

These quantifications were performed using IMOD. The long axis was identified for each of the mitochondria and mitochondrial width (diameter of the OMM; perpendicular to the long axis) was measured at different positions along the long axis. The central tomogram section for each of these positions along the mitochondria was used for this purpose and distances were measured center-to-center. Using the same central section, two measurements were made for the intermembrane distance (one on either side of the mitochondrion) at each of the positions mentioned above. These mitochondrial width measurements were plotted against their corresponding intermembrane distances (average of the two values) from a total of 15 mitochondrial constriction profiles in the boxplot-cum-beeswarm plot shown in Fig. 6C.

## Acknowledgements

We thank S. Chen and A. Malyutin at the California Institute of Technology cryo-EM facility and Bill Anderson at The Scripps Research Institute electron microscopy facility for microscope support, and Jean-Christophe Ducom at The Scripps Research Institute for computational support. We thank A. Collazo and S. Wilbert for technical assistance with confocal microscopy. We thank Catherine Oikonomou for her critical input on the manuscript. The confocal imaging was performed at the Biological Imaging Facility at Caltech, and the cryo-EM imaging was performed at the Beckman Institute Resource Center for Transmission Electron Microscopy at Caltech and the Scripps Research Institute Hazen Cryo-EM Microscopy Suite.

## Author Contribution Statement

**Mageswaran, Shrawan Kumar:** conceptualization, methodology, sample preparation, data collection, segmentation, data analysis, software/code development, figure generation, writing, editing

**Grotjahn, Danielle Ann:** conceptualization, methodology, sample preparation, data collection, segmentation, data analysis, figure generation, writing, editing, supervising

**Zeng, Xiangrui:** data analysis, software/code development, figure generation, writing, editing

**Barad, Benjamin Asher:** data analysis, software/code development, figure generation, editing

**Medina, Michaela:** sample preparation, data collection, data analysis

**Hoang, My Hanh:** writing and data analysis

**Dobro, Megan J:** segmentation

**Chang, Yi-Wei:** editing, supervising

**Xu, Min:** editing, supervising

**Yang, Wei Yuan:** conceptualization, methodology, sample preparation, data collection, editing

**Jensen, Grant J.:** conceptualization, editing, supervising

