## Supplemental Figures for "Nanoscale details of mitochondrial fission revealed by cryo-electron tomography"

### Supplementary Figures

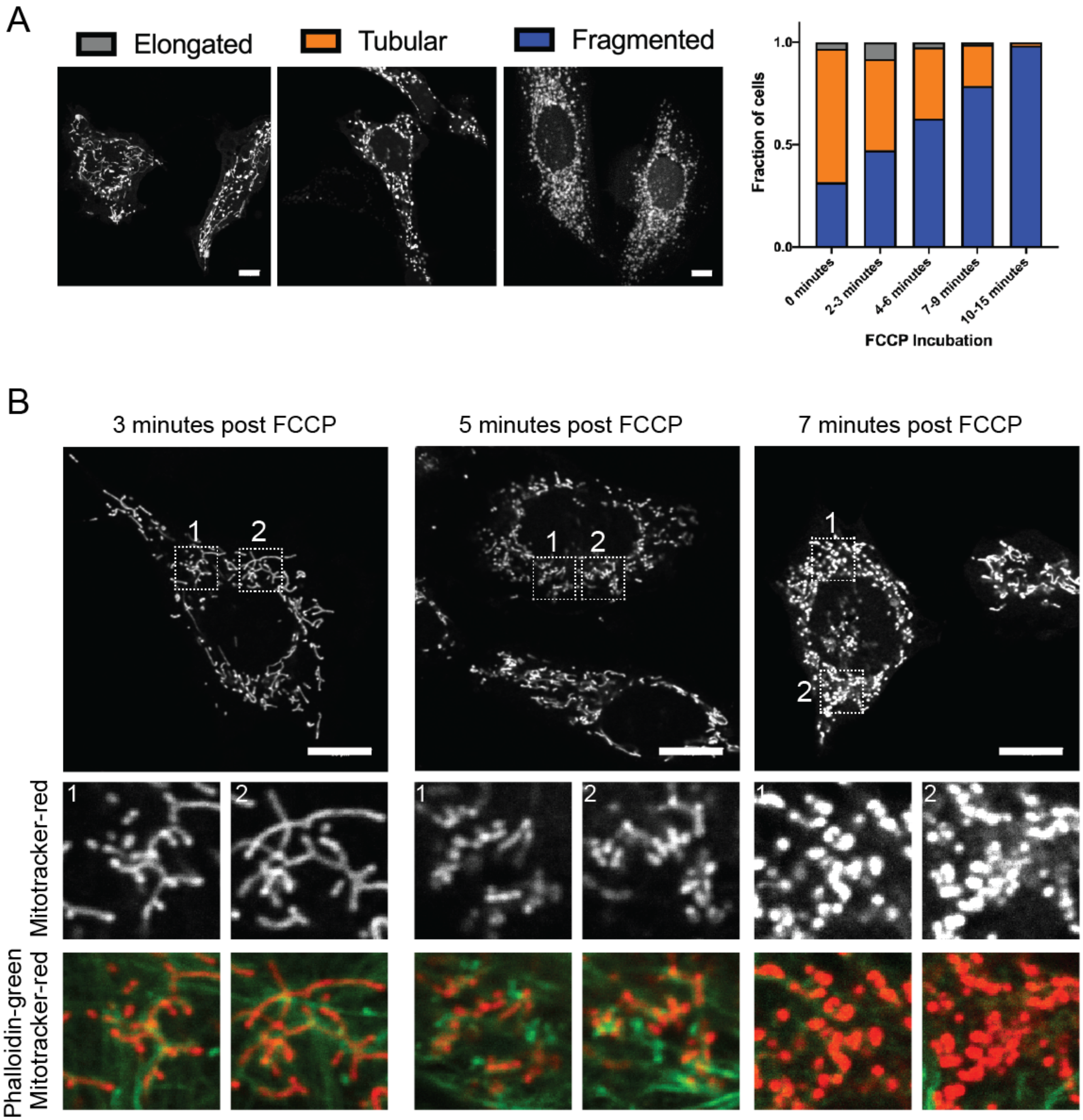


**Supplementary Figure 1. Temporal dynamics of FCCP-induced mitochondrial fission.** (**A**) Phenotypes of mitochondrial morphology (hyperfused, blue; intermediate, orange; and fragmented, gray) observed by fluorescence imaging of MitoTracker-Red-stained MEF cells following FCCP treatment. The percentage of cells exhibiting each phenotype at the indicated time after FCCP addition is shown at right. (**B**) Representative images of MitoTracker-Red-labeled cells at the indicated times after FCCP addition. Boxed regions are shown enlarged below, and overlaid with signal from phalloidin-labeled actin filaments (green). Scale bars are 10 µm.

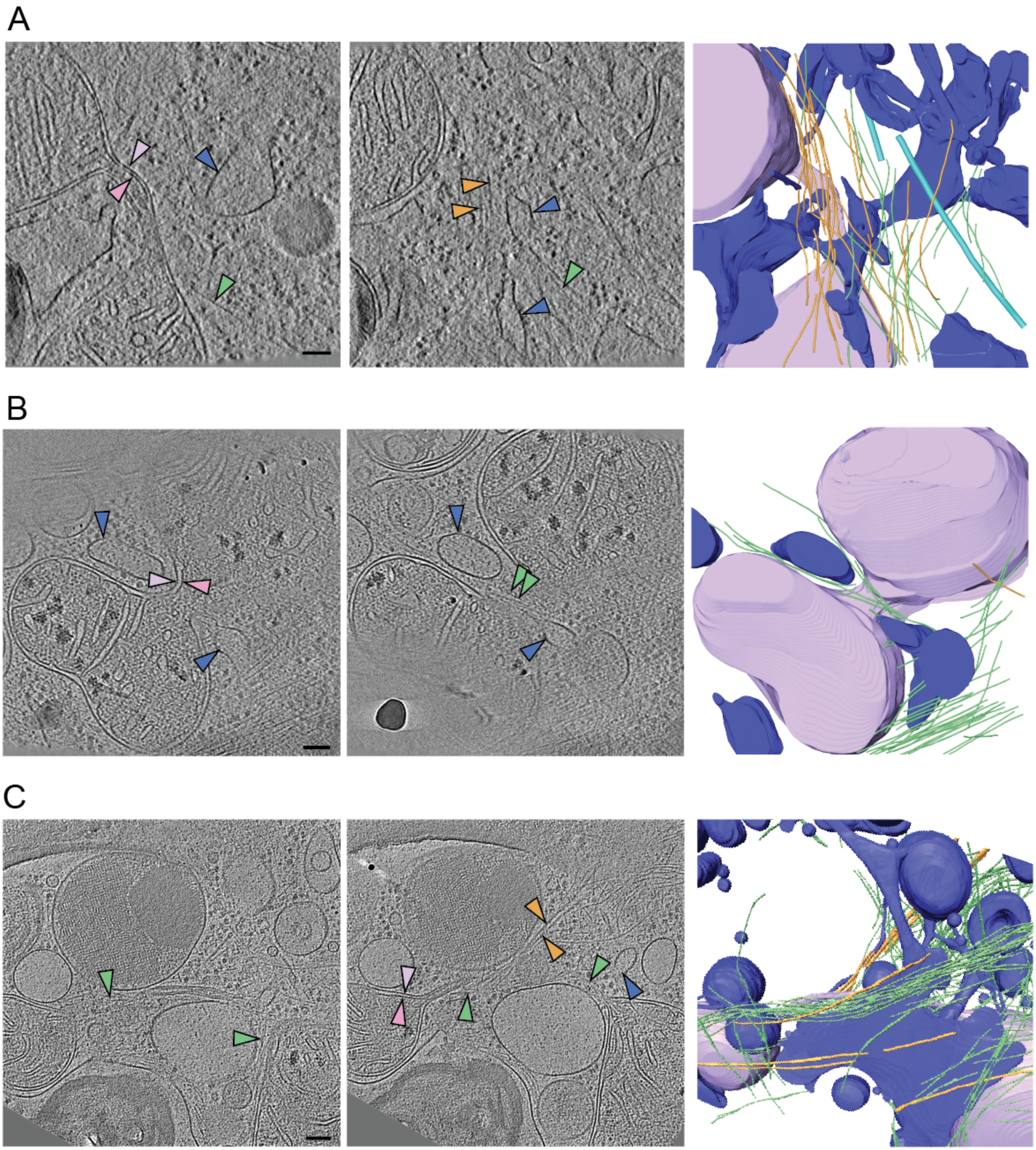


**Supplementary Figure 2. Ultrastructure of mitochondrial constriction sites is conserved across cell types, treatments.** Tomogram slices at different heights through constricting mitochondria and corresponding 3-D segmented models from (**A**) an FCCP-treated U2OS cell, (**B**) an untreated MEF cell, and (**C**) an untreated INS-1E cell. Visible features are color-coded as in other figures: OMM (purple), IMM (pink), ER (blue), F-actin (green), septin filaments (orange). Scale bars are 100 nm.
